# Response of Australian pied oystercatchers *Haematopus longirostris* to increasing abundance of the beach bivalve prey *Donax deltoides*

**DOI:** 10.1101/444786

**Authors:** Stephen L. Totterman

**Affiliations:** 179 Reedy Creek Road, Empire Vale, NSW, 2478, Australia

## Abstract

This study measured the response of Australian pied oystercatchers *Haematopus longirostris* on South Ballina Beach, New South Wales, Australia during a recovery in the stock of the primary prey species *Donax deltoides*, a large beach clam and commonly known as the ‘pipi’. It was predicted that oystercatcher counts would increase when pipi abundance increased (numerical response) and that oystercatcher feeding rates would also increase (functional response). Between Oct 2009 and Mar 2015, mean pipi density increased from c. zero to 30 pipis/m^2^. Mean oystercatcher feeding rates increased to an asymptote of c. 0.26 pipis/min. Breeding season mean counts of adult-plumage oystercatchers increased from 23 to 43, largely driven by non-territorial birds. Prey size selection was absent, both among different prey types and among pipis > 20 mm. This report provides some insights into the feeding ecology of oystercatchers on sandy ocean beaches that should be valuable in planning future studies.

## Introduction

Oystercatchers are specialist predators of large bivalves and numerous studies have investigated the close association between oystercatcher and bivalve populations (*e.g.*Goss-Custard 1996). However, the majority of those studies are from European estuaries and almost no studies of predator-prey associations for oystercatchers on sandy ocean beaches have been published (but see Ward 1991 and Taylor & Taylor 2005).

The Australian pied oystercatcher *Haematopus longirostris* inhabits estuaries and sandy ocean beaches around the Australian coast, with major populations in the south-eastern states of Tasmania, South Australia and Victoria (Taylor *et al.* 2014). The species is non-migratory and many breeding pairs remain on their all-purpose territories throughout the year.

Commonly known as the ‘pipi’, *Donax deltoides* is a large bivalve that inhabits the intertidal and shallow subtidal zones on sandy ocean beaches from Fraser Island, Queensland, to the Murray River, South Australia (Ferguson *et al.* 2014). Pipi are fast-growing and short-lived; in New South Wales (NSW), pipi recruits grow to a shell length of 37 mm within ten months, at which size 50% of pipis are sexually mature, and adult pipis continue growing to an asymptotic length of 75 mm in c. 18–22 months (Murray-Jones 1999). Pipis are the major prey for Australian pied oystercatchers on beaches in NSW. Owner and Rohweder (2003) reported that oystercatchers aggregated at beaches with high pipi biomass (high mean pipi numerical density and large mean pipi length). Pipis are also exploited by humans for food and as bait for recreational fishing. Owner and Rohweder (2003) and Harrison (2009) raised concerns about the potential negative impacts of commercial fishing on pipi stocks and oystercatcher populations.

NSW commercial pipi landings declined from a peak of 670000 kg in 2001 to 9000 kg in 2011 (Gray 2016a). Pipi density on South Ballina Beach declined from 20/m^2^ in 2003 (Harrison 2009) to c. zero in 2009 (this study). Harrison (2009) associated a decline in oystercatcher counts from northern NSW with this 2003–2009 ‘pipi crash’. Fishery management responses to the pipi crash, introduced in 2012, were an annual Dec–Jun commercial fishing closure, closures of certain beaches and zones within certain beaches, a daily trip limit of 40 kg per commercial fisher and a minimum legal length limit of 45 mm (Ferguson *et al.* 2014; Gray 2016a). Only hand gathering of pipis is allowed in NSW.

Assuming the 2003–2009 pipi crash was temporary, it was predicted that oystercatcher counts and feeding rates would increase when pipi abundance increased. This study measured the recovery in the pipi stock and responses of oystercatchers on South Ballina Beach between 2009–2015.

## Materials and methods

### Study site

South Ballina Beach (Fig. 1) is 23 km long between Broadwater Headland (29°02’16”S, 153°27’12”E) and the Richmond River (28°52’35”S, 153°35’08”E). It is an intermediate energy, wave-dominated beach with a longshore bar and trough outer bar and transverse bar and rip inner bars (Short 2007). The intertidal zone is gently sloping with fine-to medium-grained sand (0.125–0.5 mm). Fore dune vegetation is dominated by hairy spinifex *Spinifex seriseus*. The dune and hind dune are dominated by bitou bush *Chrysanthemoides monilifera* and coastal wattle *Acacia longifolia sophorae*. Adjoining land-use is mostly sugar cane farming for the central 17 km with protected areas in the south (Broadwater National Park) and in the north (Richmond River Nature Reserve). Human population densities are low and scattered except for the villages of South Ballina in the north (17 houses plus one holiday park) and Patch’s Beach in the middle (18 houses).

**Fig. 1.**
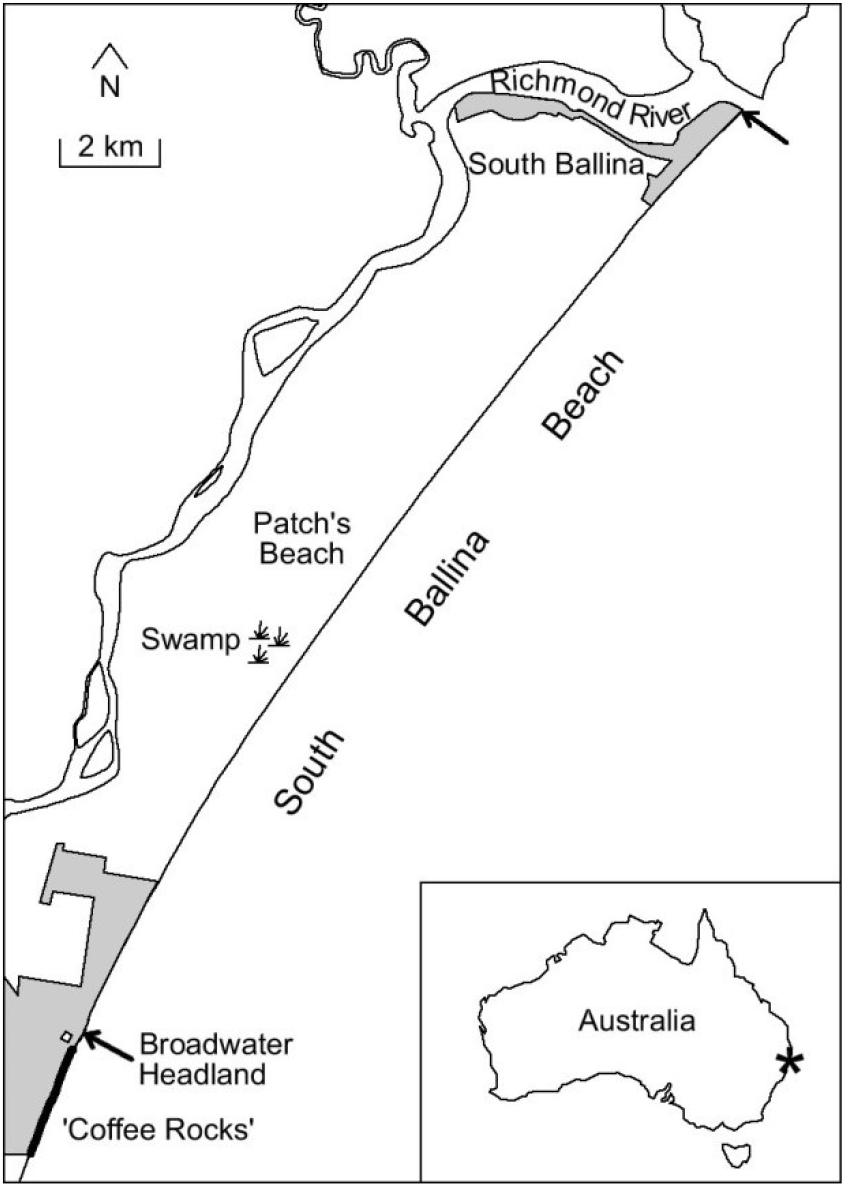
Map of South Ballina Beach and inset map showing location within Australia. Arrows indicate the southern and northern boundaries of the study site. Oystercatcher counts included the ‘coffee rocks’, which was used by small numbers of non-territorial oystercatchers as a roost site and the swamp, which was used by some oystercatchers as a refuge during and soon after bad weather. Grey areas are Broadwater National Park (south) and Richmond River Nature Reserve (north).

South Ballina Beach has the largest breeding population of any single beach in NSW (it is rare to find more than one or two breeding pairs on other beaches along the NSW coast except for the ‘Bombing Range’ end of Ten Mile Beach; unpub. data). The local breeding season is August– December and late attempts generally are replacement clutches (Wellman *et al*. 2000). South Ballina is also a major pipi fishery (Owner 1997). Commercial pipi harvesting on South Ballina Beach stopped in 2004 (Harrison 2009) and resumed in 2013, during this study. Harvesting was restricted to the southern 12 km of the beach and annually between 1 Jul and 30 Nov. There were c. six active commercial fishers on the beach from 2013 (Gray 2016a), similar to historical numbers (pers. obs.).

Field studies started in Oct 2009 and ended in Feb-Mar 2015. Collection of field data was biannual: in the pre-breeding/breeding months (Jul–Oct; hereafter ‘breeding season’) and in the post-breeding months (Jan–Mar; hereafter ‘non-breeding season’). The earliest observations of copulating oystercatchers were from Jul.

### Pipi abundance

Pipis typically aggregate in ‘bands’ that run parallel to the shoreline. Previous use of the term ‘pipi beds’ (Owner & Rohweder 2003; Harrison 2009) is confusing because aggregations of pipis are not static like the cockle and mussel beds in numerous studies for Eurasian oystercatchers *Haematopus ostralegus*.

Pipis were sampled at low tide by digging and sieving square 0.1 m^2^ × 20 cm deep quadrats (James & Fairweather 1995; Harrison 2009). Across shore transects were used to locate aggregations of pipis and then seven replicate quadrats were sampled within the pipi band (Owner & Rohweder 2003; Harrison 2009). Four pipi band quadrats were sampled along a c. 45° diagonal immediately north of the transect, with five paces between quadrats, and three quadrats immediately south. If a transect did not locate an aggregation then pipi band sampling was performed at the same levels as pipi bands in adjacent transects. Closely-spaced quadrats are pseudoreplicates (Millar & Anderson 2004) and so pipi density was calculated as the mean number of pipis per transect pipi band (divided by 0.1 m^2^ × 7 = 0.7 m^2^ to get pipis/m^2^). The sampling unit is the transect pipi band and mean pipi density was measured at the beach scale (*i.e.* the mean of replicate transect pipi bands). Five to 10 transects were sampled on each biannual sampling occasion, equally-spaced (according to the relevant transect sample size) along the beach and starting from a random distance (*i.e.* systematic sampling with a random start).

Sieve mesh size was 6 × 6 mm although ‘finger-raking’ of flooded quadrats (James & Fairweather 1995) was often used in the swash zone, where thixotropic sand and frequent waves make excavation of quadrats difficult. Pipis retained in the sieve were measured with a calliper to the nearest mm (maximum shell length). Pipis smaller than ten mm passed diagonally through the sieve and were not counted. Murray-Jones (1999) classified pipis < 16 mm as new recruits. Very small bivalves are often ignored in oystercatcher feeding studies because they are swallowed whole with practically zero handling time and are often unidentified in feeding observations, biomass increases exponentially with shell length and small bivalves contribute little to ‘intake rates’ (*i.e.* the rate at which prey biomass or energy is consumed; Goss-Custard *et al.* 2006) and, in some studies, the birds ignore new recruits (*e.g.* Ens & Goss-Custard 1984; Ens *et al.* 1996b).

### Oystercatcher counts

Oystercatchers were counted from a four-wheel drive vehicle driven along South Ballina Beach at about 40 km/h, stopping briefly to count birds, check for leg bands and flags and record the distance from the Boundary Creek four-wheel-drive access track (zero km). Beach driving was unrestricted on South Ballina Beach and the birds are habituated to slow moving vehicles. The ‘coffee rocks’ south of Broadwater Headland were surveyed on foot when erosion prevented driving. An intermittent swamp near the middle of the beach was surveyed using binoculars and a 20–60× telescope from the adjacent dunes (Fig. 1).

‘Adult’-plumage oystercatchers were counted including second immature plumage attained at c. one year old, which looks the same without close inspection (Marchant & Higgins 1993). One and two year old oystercatchers have relatively dull colours of bill, legs and iris compared to adults (Marchant & Higgins 1993), but the 23 km long beach had to be surveyed quickly to avoid movements of birds confounding the counts and not all individuals could be inspected closely (*e.g.* birds at distance, in unfavourable light or flying past).

Territory mapping (spot mapping) was applied from Jul 2012, in both breeding and non-breeding seasons. Three counts were made on each biannual sampling occasion at intervals of 5–16 days. Oystercatcher territories were identified as adult ‘pairs’ that were present on the same stretch of beach on at least two of three counts. Three counts was considered sufficient for a low-cost estimate of oystercatcher territories because oystercatchers on the beach are conspicuous and easy to find, territories are one-dimensional and several territories are well known from field studies starting in the 1990s (Wellman *et al.* 2000). Interpretation of spot mapping results was aided by observations of leg bands and flags on some birds and observations of territorial and breeding behaviours (especially when watching focal birds for feeding observations, see Feeding observations below). However, it must be noted that these territory estimates are short-term results using standardised methods and not all ‘pairs’ that occupy a stretch of beach early in the breeding season persist and go on to breed (*i.e.* mapped territories > breeding pairs). Counts of breeding pairs are costly, requiring frequent visits throughout the breeding season.

With knowledge of territories, total counts could be separated into territorial adults and non-territorial birds (non-territorial count = total count – territorial birds present during the count). Before Jul 2012, a local expert assisted with the identification of oystercatcher territories (B.G. Totterman, pers. comms.).

### Feeding observations

Foraging effort of oystercatchers on sandy ocean beaches is spatially and temporally irregular. Pipis undergo tidal migrations up and down the beach face and oystercatchers can feed in the swash zone through most of the tidal cycle (Owner 1997). Aggregations of sandy beach bivalves are spatially variable and foraging oystercatchers often travel > 100 m along the shoreline, presumably searching for aggregations (Taylor & Taylor 2005). Oystercatchers can also consume large bivalves faster than they can digest them (Kersten & Visser 1996). This ‘digestive bottleneck’ plus the extended availability of prey can explain the general ‘laziness’ observed for oystercatchers on sandy ocean beaches, with long intervals between short feeding sessions (Taylor & Taylor 2005). An observer needs to be patient, waiting between oystercatcher feeding sessions or opportunistic, searching the beach for feeding birds.

Focal animal sampling was used in this study and swash zone feeding observations were mostly from territorial oystercatchers (*i.e.* patient strategy). These were more reliably located than were non-territorial birds. Breeding season observations avoided active breeders with unfavourable time-budgets. Sample sizes were increased with observations from adult-plumage non-territorial oystercatchers (*i.e.* opportunistic strategy). Observations were free from interference from conspecifics because no other oystercatchers were in close proximity to the selected focal bird. Territorial pairs do often feed simultaneously but usually some tens of metres apart (pers. obs.). Non-territorial oystercatchers selected were alone or apart from the flock. Interference from gulls was rare, unlike Taylor and Taylor (2005), because large gulls are not locally resident and silver gulls *Chroicocephalus novaehollandiae* were not abundant. Selection of oystercatchers for feeding observations was not random but care was taken to select different birds (territories) along the beach.

Feeding observations were made during ebb tides (between 1–4 hrs after high tide). Birds were observed with a 20–60× telescope. Individual feeding observations lasted from the start of a feeding session until the bird stopped feeding or until a maximum of 20 min, to prevent observer fatigue. Data were recorded with a ten s sampling interval and so each activity record represents a ten s ‘block’. Activities were classified into five major categories: moving (walk, run, fly), standing, maintenance (scratch, preen, bathe), foraging (search, peck, sew, probe, transport, handle, eat) and interspecific (sky watching, attack kleptoparasite). Feeding behaviours are as described in Hulscher (1996), except that Australian pied oystercatchers are not known to attack bivalves by ‘hammering’ into the shell (Marchant & Higgins 1993; Taylor & Taylor 2005). Prey eaten were counted in five categories: pipi, polychaete worm, crab, fish and unidentified small prey. Small prey were often unidentified because they were swallowed whole and the birds were observed from some distance. Individual pipi feeding rates were calculated as the sum of pipis eaten divided by the active foraging time (*i.e.* the sum of search, peck, sew, probe, transport, handle and eat activities × 10 s; Goss-Custard *et al.* 2006). Very short feeding sessions < 2 min duration can provide inaccurate feeding rates (*e.g.* dividing by small numbers inflates results) and were discarded (Ens & Goss-Custard 1984; Ens *et al.* 1996a). The sampling unit is the individual feeding observation and mean feeding rate was measured at the beach scale (*i.e.* the mean of feeding observations along the beach, within each biannual sampling period).

### Shell sampling

Oystercatchers feeding on large bivalves leave behind spent shells and these can sometimes be measured to estimate the sizes of consumed prey (Sutherland 1982; Ward 1991; Owner 1997; Taylor & Taylor 2005). The specialised feeding methods of oystercatchers result in specific ‘tell-tale’ characters on depredated bivalves (Sutherland 1982; Ward 1991). Australian pied oystercatchers attack buried pipis by stabbing between the valves at the upward-facing, anterior end and severing the adductor muscles (Marchant & Higgins 1993). Shells left by oystercatchers are usually complete, with the two valves forming an acute angle. The valves are clean, with only shreds of adductor muscle attached. The valves may be slightly chipped at the posterior end, resulting from failed stabbing attacks. Pipi shells opened by recreational fishers for bait are nearly always smashed and the valves are not cleaned of all flesh. Stranded pipis are sometimes found above the mean high tide drift line. The valves of stranded pipis are initially closed and the dead animal becomes desiccated with exposure.

Pipi shells were sampled from ten 500 m sections of South Ballina Beach, randomly selected, without replacement on each biannual sampling occasion. Shells were collected on ebb and low tides by searching between the toe of the fore dune and the top of the swash zone. Two or three passes were required to cover the beach width, which increased as the tide receded. Pipi shells were measured with a calliper to the nearest mm (maximum shell length). Shells were then discarded off the beach, so that they would not be collected again. Length-frequency distributions were measured at the beach scale, by pooling together all measurements within each biannual sampling period.

### Statistical analyses

All biannual data were averaged at the beach scale. This scale of investigation is very large compared to sampling plots in European oystercatcher studies, however the same logic applies: mean oystercatcher counts and feeding rates were correlated with mean prey abundance for matched sampling occasions.

All statistical analyses were performed using R version 3.4.4 (R Core Team 2018). Numerical and functional response models were fitted using least squares regression. Repeated data collection over time often leads to temporal autocorrelation, where a result at some point in time is related to, and not independent of, nearby values. Least squares regression coefficients are unbiased in the presence of autocorrelated residuals, but the standard errors of these coefficients are not reliable (Allen 1997). Autocorrelation was investigated with partial autocorrelation plots and Durbin-Watson tests were computed using *lmtest* version 0.9-36 (Zeileis & Hothorn 2002). Time series intervals were slightly uneven (mean six months, range 5–8 months), however biannual sampling occurred during oystercatcher breeding or non-breeding seasons, where oystercatcher counts and territory occupancy are reasonably stable. Pipi density, excluding new recruits, is also not expected to show large changes from month to month. Sets of candidate regression models were compared using small sample corrected Akaike’s information criterion (*AICc*) and Akaike weights computed using *AICcmodavg* version 2.1-1 (Mazerolle 2017). Akaike weights sum to one for the entire set of candidate models and can be interpreted as the weight of evidence in favour of a given model (Johnson & Omland 2004).

The asymptotic hyperbolic function is generally assumed for shorebird functional responses (Goss-Custard *et al.* 2006):

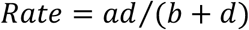

where *a* is the asymptote and *d* is the numerical prey density. The ‘half-asymptote density’ *b* is the prey density at which the rate has risen to half of the asymptote. A zero half-asymptote (*b* = 0) gives a constant rate function: *Rate* = *a*.

Pipi sampling included the 2013 and 2014 fishing seasons and fished and non-fished management zones. Separate before-after, control-impact (BACI) analyses were performed for the before- and during-fishing samples in 2013 and 2014 (similar to Gray 2016b). After-fishing samples were not included because those data were strongly affected by growth of smaller pipis. The BACI model used is:

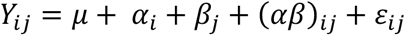

where *Y_ij_* is the observation, *μ* is the overall mean, *α_i_* is the effect of period (*i* = before/during fishing), *β_j_* is the effect of location (*j* = fished/non-fished zone), (*αβ*)_*ij*_is the interaction between period and location and *ε_ij_* represents residual error (Smith 2002). A significant interaction indicates a different response between zones that could be interpreted as a fishing effect. BACI pipi counts (integerized pipi band densities) were modelled with negative binomial generalised linear models computed using the MASS package (Venables & Ripley 2002). BACI pipi length sample sizes were large and those data were modelled with ordinary least squares models. Analysis of deviance/variance tables with type III sums of squares for unbalanced designs were computed using *car* version 3.0.0 (Fox & Weisberg 2011). Temporal autocorrelation is not relevant for these two-sample BACI models because there was no temporal replication in the before- and during-fishing periods. Spatial autocorrelation could be present but transects were well spaced at 2.5 km apart in Jan 2013 and 2 km apart on subsequent sampling occasions. Small transect sample sizes were assumed to have the greatest effect on the precision of density estimates.

## Results

### Pipi abundance

Mean pipi density increased from c. zero to 31/m^2^ during this study (Fig. 2A). Large variation among transects within each biannual sampling occasion indicates strongly aggregated along shore distributions of pipis. Peaks in Jan 2013 and Feb 2015 followed recruitment events in Jul 2012 and Sep 2014 respectively (Fig. 3). This time lag occurred because pipis < 10 mm were not sampled and perhaps also because recruitment continued for a number of months. Apart from those two recruitment events, < 10% of pipis measured were ≤ 20 mm and pipi > 20 mm densities (Fig. 2B) were similar to total densities. Further analyses concern pipi > 20 mm densities (see pipi size selection results below) although results using total densities were similar.

**Fig. 2.**
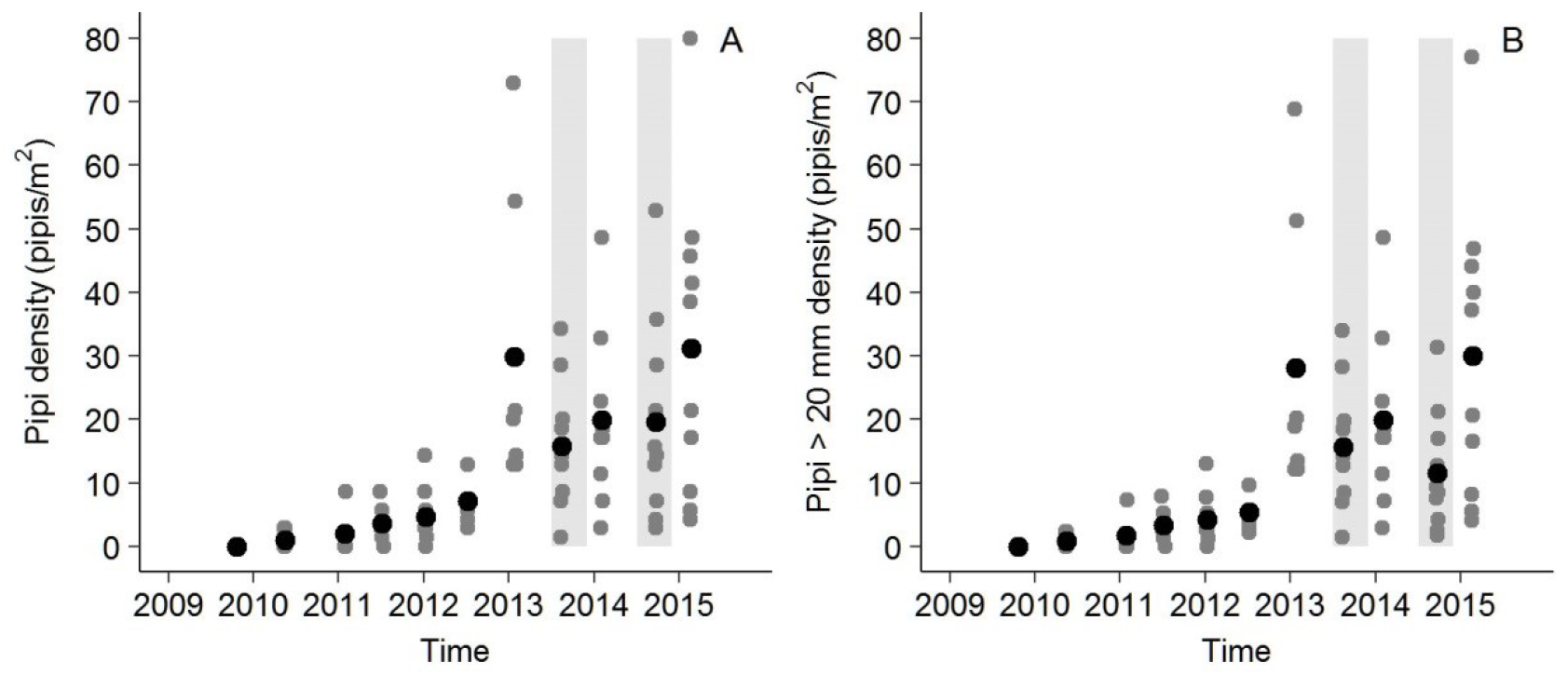
Pipi total density (A) and pipi > 20 mm density (B) over time at South Ballina Beach. Black-filled circles are mean densities. The time scale is continuous, with tick marks at 1 Jan. Light-grey vertical bands indicate commercial shellfishing seasons. Pipi > 20 mm densities were used for subsequent analyses.

**Fig. 3.**
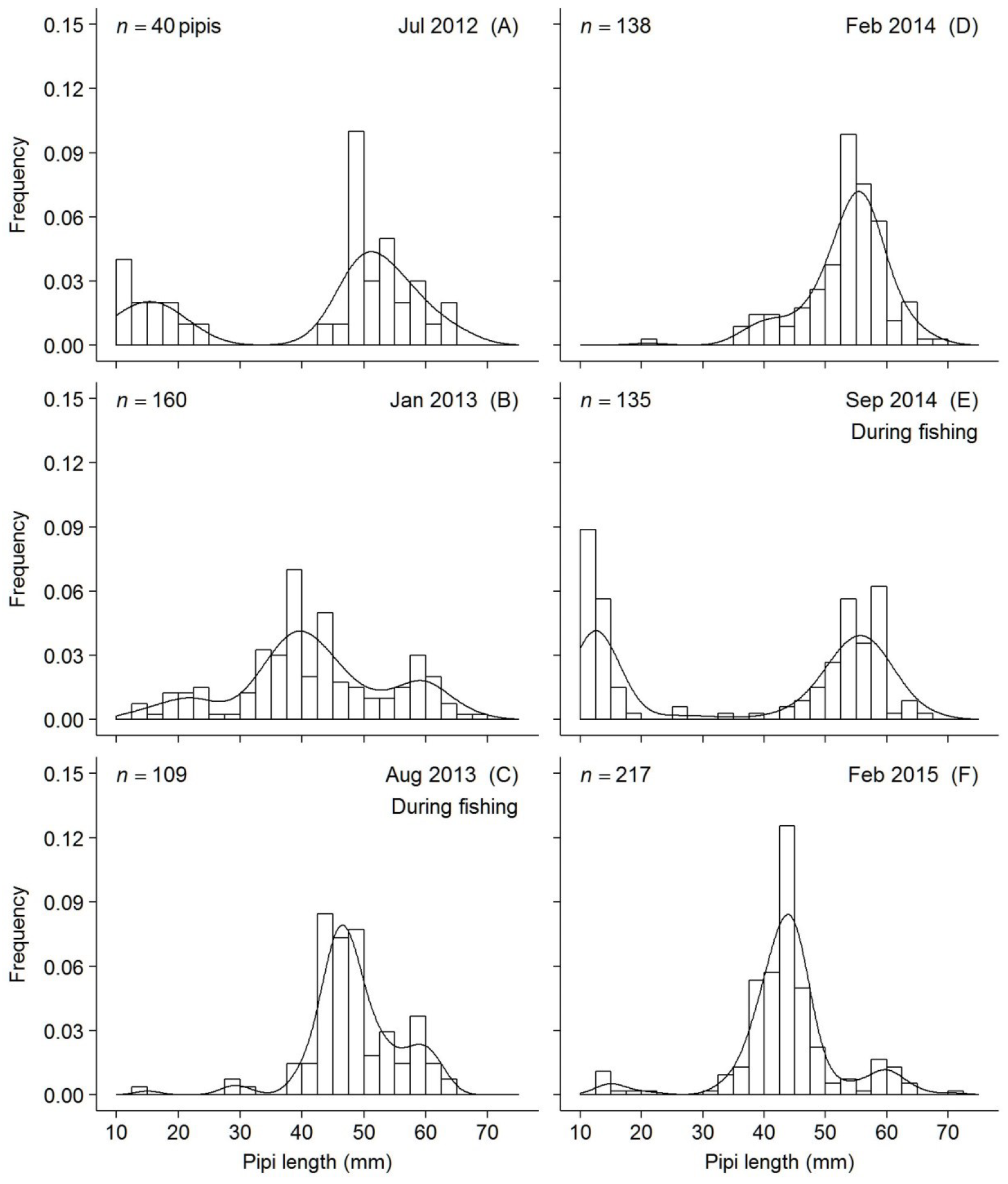
Length-frequency distributions for live pipis at South Ballina Beach (for samples with *n* ≥ 30 pipis). Lines are kernel densities. Bimodal distributions from Jul 2012 (A) and Sep 2014 (E) indicate recruitment events. There was commercial shellfishing in 2013 and 2014 (C & E).

Commercial shellfishing resumed in Jul 2013 and c. 10400 kg of pipis were harvested from South Ballina Beach in that year (Gray 2016b). Catch data are not available for 2014. Beach scale pipi > 20 mm mean density decreased during fishing by 42% in 2013 and 44% in 2014 (Fig. 2B). Analysis of deviance (negative binomial generalised linear models) indicated only beach scale changes in pipi density and no differences between unfished and fished zones (Table 1, Fig. 4A). Pipi > 20 mm mean length comparisons were affected by growth (Fig. 3). For 2013, ANOVA indicated consistently larger pipi > 20 mm mean lengths in the non-fished zone and continued growth of the Jul 2012 cohort (Table 2, Fig. 4B). For 2014, ANOVA indicated a decrease in mean length in the unfished zone but an increase in the fished zone (*P* < 0.001), which is opposite to the expected fishing impact (Table 2, Fig. 4B).

**Table 1.**
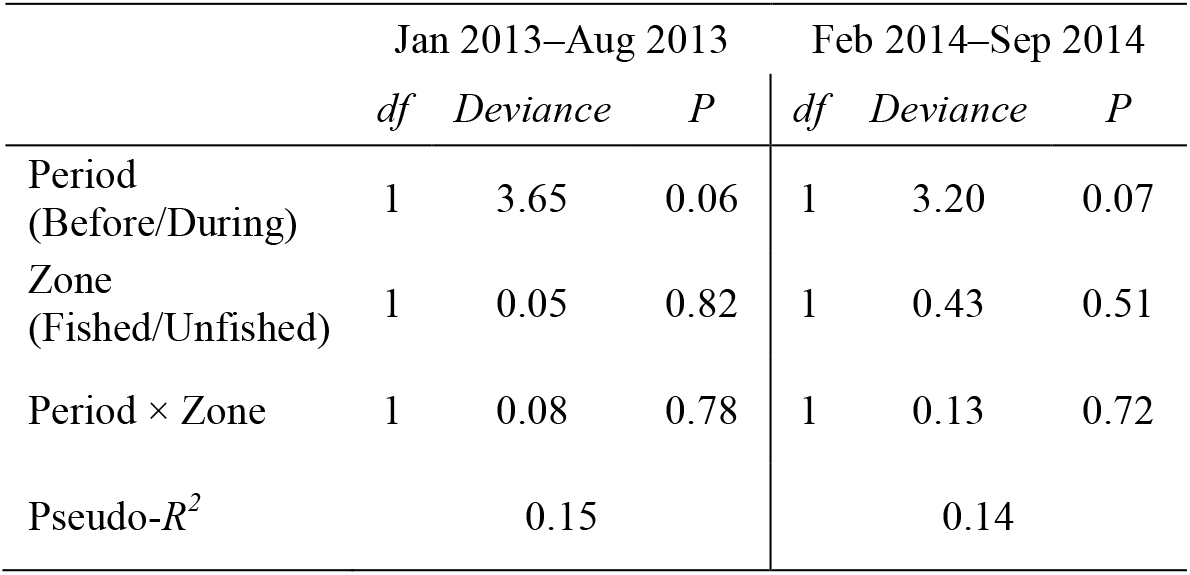
Pipi > 20 mm density analysis of deviance (negative binomial generalised linear model) results for the 2013 and 2014 fishing seasons. A large Period × Zone interaction deviance statistic indicates a different response between zones. See Fig. 4A for mean densities. Pseudo-*R*^2^ is simply the squared Pearson’s product-moment linear correlation between observed and fitted values.

|  | Jan 2013–Aug 2013 |  |  | Feb 2014–Sep 2014 |  |  |
| --- | --- | --- | --- | --- | --- | --- |
|  | *df* | *Deviance* | *P* | *df* | *Deviance* | *P* |
| Period<br>(Before/During) | 1 | 3.65 | 0.06 | 1 | 3.20 | 0.07 |
| Zone<br>(Fished/Unfished) | 1 | 0.05 | 0.82 | 1 | 0.43 | 0.51 |
| Period × Zone | 1 | 0.08 | 0.78 | 1 | 0.13 | 0.72 |
| Pseudo-$R^2$ | | 0.15 | | | 0.14 | |

**Table 2.** Pipi > 20 mm length ANOVA (linear model) results for the 2013 and 2014 fishing seasons. A large Period × Zone interaction *F* statistic indicates a different response between zones. See Fig. 4B for mean lengths.

|  | Jan 2013–Aug 2013 |  |  |  | Feb 2014–Sep 2014 |  |  |  |
| --- | --- | --- | --- | --- | --- | --- | --- | --- |
|  | *df* | *SS* | *F* | *P* | *df* | *SS* | *F* | *P* |
| Period<br>(Before/During) | 1 | 1883 | 21.8 | <0.001 | 1 | 4 | 0.1 | 0.75 |
| Zone<br>(Fished/Unfished) | 1 | 760 | 8.8 | 0.003 | 1 | 733 | 17.0 | <0.001 |
| Period × Zone | 1 | 131 | 1.51 | 0.22 | 1 | 723 | 16.8 | <0.001 |
| Residuals | 255 | 22023 |  |  | 260 | 11217 |  |  |
| $R^2$ | | | 0.11 | | | | 0.11 | |

**Fig. 4.**
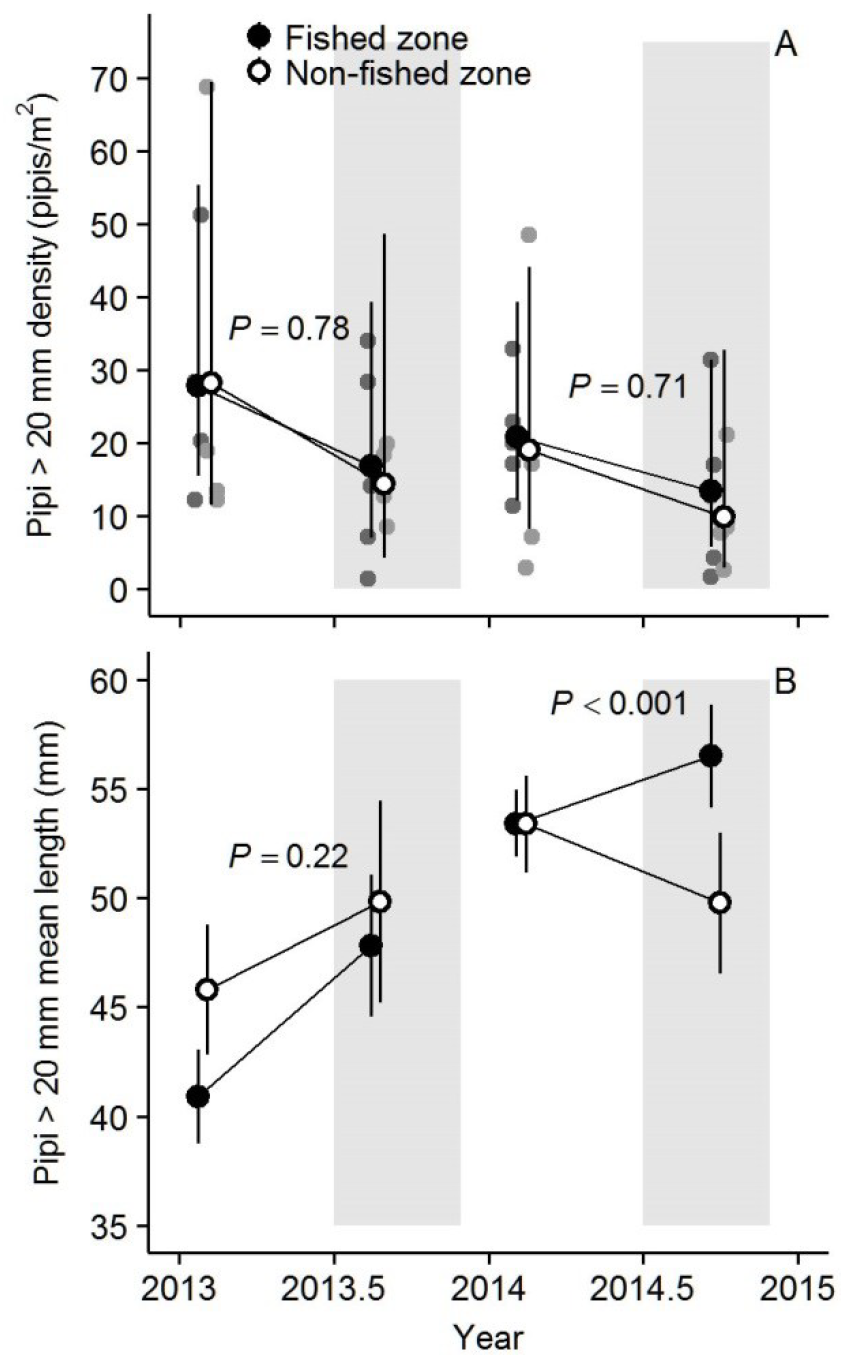
Changes in pipi > 20 mm mean densities (A) and lengths (B) in fished (black fill) and non-fished (white fill) zones on South Ballina Beach for the 2013 and 2014 fishing seasons (light-grey areas). *P*-values refer to the interaction period (before/during fishing) × zone (Tables 1 & 2). Results for the non-fished zone are shifted slightly right to avoid over plotting.

### Diet and prey size selection

Australian pied oystercatchers swallow bivalves smaller than c. 20 mm whole (Evans 1975; Taylor & Taylor 2005) and the minimum pipi length from shell samples was 24 mm. Pipis > c. 20 mm were 56% of 241 prey items eaten by oystercatchers during swash zone feeding observations, unidentified small prey were 41% and polychaete worms were 2%. Some of the 41% unidentified small prey could have been small pipis, particularly during the recruitment events in Jul 2012 and Sep 2014. Length-frequency distributions for pipis taken by oystercatchers (from shell samples) were similar to those for pipis > 20 mm available on the beach (from quadrat samples; Fig. 5). Across these comparisons, paired mean differences were small and showed no consistent positive or negative bias. A least squares regression estimate for the overall paired mean difference was 1.0 mm (shells > live pipis; 95% CI −1.0 to 3.0). There was no significant autocorrelation in the residuals (*ρ*_1_ = −0.39, *DW* = 2.16, *P* = 0.41) and this linear regression model is equivalent to a paired *t*-test.

**Fig. 5.**
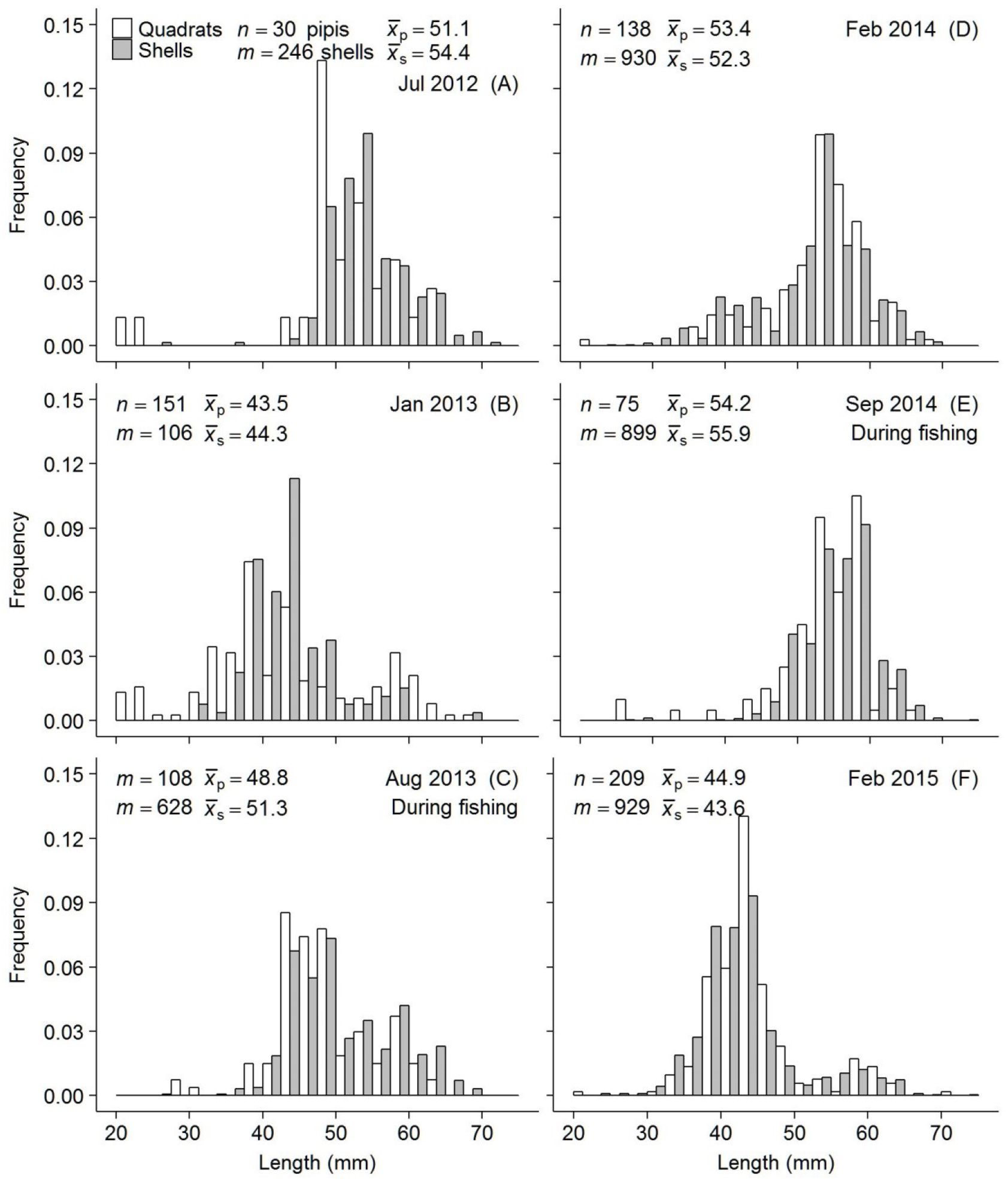
Length-frequency distributions for live pipis (from quadrat samples, grey fill) and pipis taken by oystercatchers (from shell samples, white fill) at South Ballina Beach (for samples with *n* ≥ 30 pipis). Bivalves smaller than 20 mm are swallowed whole (Evans 1975, Taylor & Taylor 2005) and have been removed from quadrat samples for these comparisons. There was commercial shellfishing in 2013 and 2014 (C & E).

### Numerical response

Total oystercatcher counts increased strongly with pipi > 20 mm density (Fig. 6A). Narrow count ranges suggest that results from Jul 2012 onwards were not sensitive to increased survey effort (three counts versus one or two previously). The numerical response was largely driven by non-territorial adult-plumage birds, for which counts increased linearly with pipi density (Fig. 6B). Three models for non-territorial counts were compared: 1) linear response (*AICc* weight = 0.85), 2) linear response with breeding/non-breeding season covariate for the slope (*AICc* weight = 0.15), and 3) linear response model with season covariates for both the slope and intercept (*AICc* weight = 0.01). The ordinary least-squares slope for non-territorial oystercatcher counts model one (Fig. 6B2) was 1.64 (95% CI = 1.08–2.20). The intercept was 6.2 and the accompanying 95% confidence interval −2.4 to 14.7 suggests that non-territorial oystercatcher counts are approximately zero when pipi density is zero. There was no significant autocorrelation in the residuals (*ρ*_1_ = 0.17, *DW* = 1.23, *P* = 0.96). The final three years of counts showed non-breeding season peaks, for which non-territorial oystercatcher counts model two is a better fit. Observations of leg flags indicated that these non-breeding season counts included some birds that were not present in the preceding breeding season and therefore from outside South Ballina Beach.

**Fig. 6.**
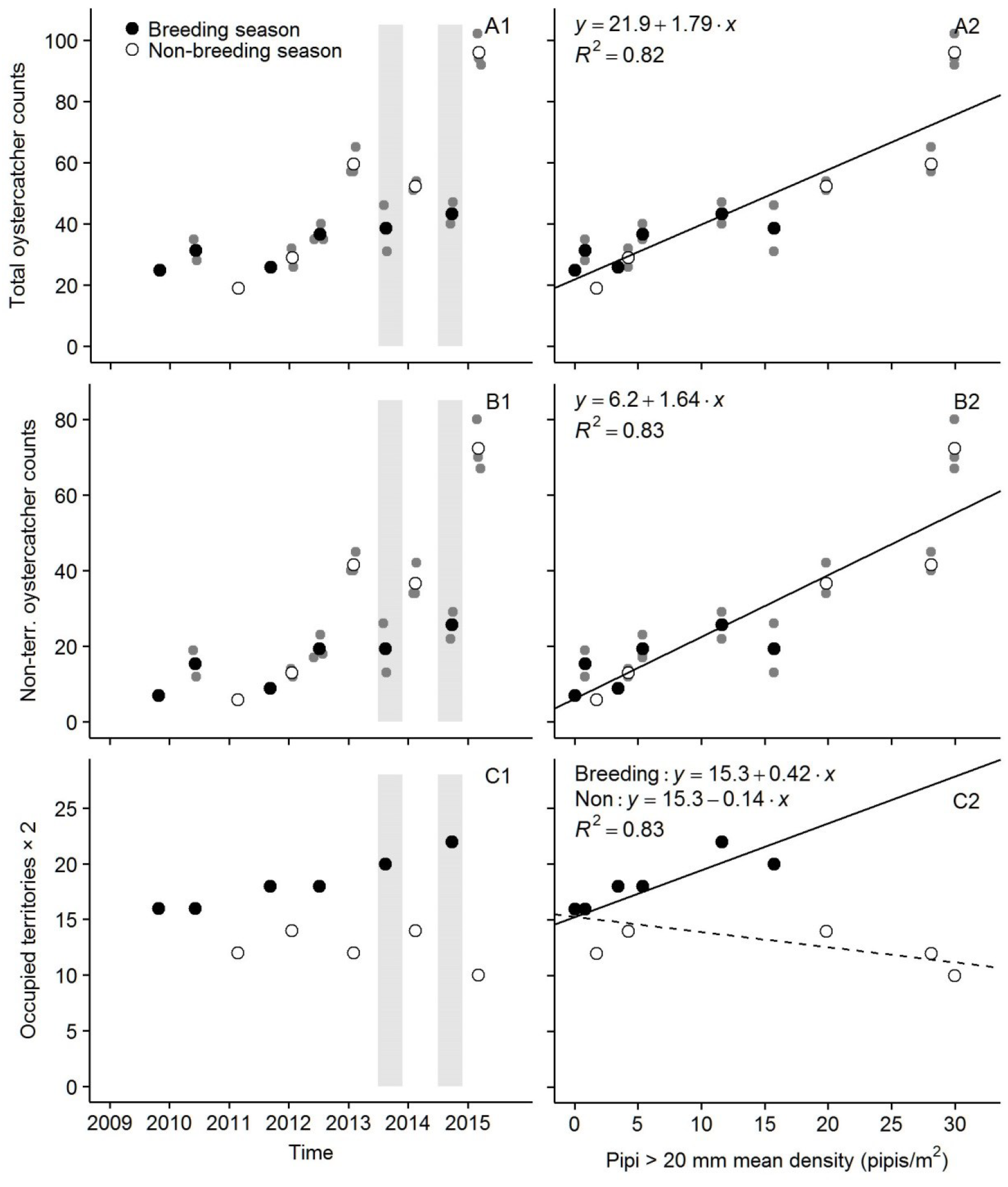
Counts of all oystercatchers (A), non-territorial, adult-plumage oystercatchers (B) and territorial oystercatchers (C) at South Ballina Beach over time (1) and numerical response to increasing pipi density (2). Black-filled circles are breeding season means and white-filled circles are non-breeding season means. Territory mapping results are point estimates. The time scales (1) are continuous, with tick marks at 1 Jan. Light-grey vertical bands (1) indicate commercial shellfishing seasons. Linear models (2) were fitted by non-linear least-squares regression and the territorial oystercatchers model (C2) includes season as a covariate.

Mapped oystercatcher territories (*i.e.* pairs) were multiplied two for regression analyses so that the numerical response can be compared to the non-territorial oystercatcher results. Territorial oystercatcher counts increased linearly with pipi > 20 mm density and there was strong seasonal variation, with fewer territories occupied in the non-breeding season (Fig. 6C). Observations of leg flags confirmed that some oystercatchers were ‘off-territory’ and counted together with non-territorial birds in non-breeding seasons. Two models for territorial oystercatcher counts were compared: 1) linear response with breeding/non-breeding season covariate for the slope (*AICc* weight = 0.77), and 2) linear response model with season covariates for both the slope and intercept (*AICc* weight = 0.23). The ordinary least-squares breeding season slope for territorial oystercatcher counts model one (Fig. 6C2) was 0.42 (95% CI = 0.17–0.67) and the non-breeding slope was −0.14 (95% CI = −0.36 to 0.09). There was no significant autocorrelation in the residuals (ρ_1_ = −0.39, *DW* = 2.70, *P* = 0.10). The substantial intercept of 15.3 individuals (*i.e.* 7–8 pairs; 95% CI = 13.4–17.1 individuals) shows that not all territories were abandoned during the 2003–2009 pipi crash.

### Functional response

Oystercatcher mean feeding rates increased with pipi > 20 mm density (Fig. 7). Individual feeding rates were highly variable, like individual pipi band densities but the modest correlation between coefficients of variation for these two variables (*r* = 0.37 or 0.09 without one high *CV* influential point) indicates that variability in feeding rates was not simply following alongshore variability in pipi abundance. Four functional response models were compared: 1) constant feeding rate (*AICc* weight = 0.15), 2) constant feeding rate with breeding/non-breeding season covariate for the intercept (*AICc* weight = 0.05), 3) asymptotic feeding rate (‘type II’ functional response, *AICc* weight = 0.80), and 4) asymptotic feeding rate with season covariate for the asymptote (*AICc* weight = 0.00). It is obvious from the functional response scatterplot (Fig. 7B) that a linear feeding rate with zero intercept model (‘type I’ functional response) does not fit. The non-linear least squares asymptote for feeding rate model three was 0.26 pipis per min of active foraging time (95% CI = 0.18–0.36). The half-asymptote density was 0.5 pipis/m^2^ and the accompanying 95% confidence interval −0.24 to 3.8 suggests a constant feeding rate (feeding rate model one). More data near zero prey density are required for precise estimates of the half-asymptote. There was no significant autocorrelation in the residuals (ρ_1_ = −0.45, *DW* = 2.70, *P* = 0.11).

**Fig. 7.**
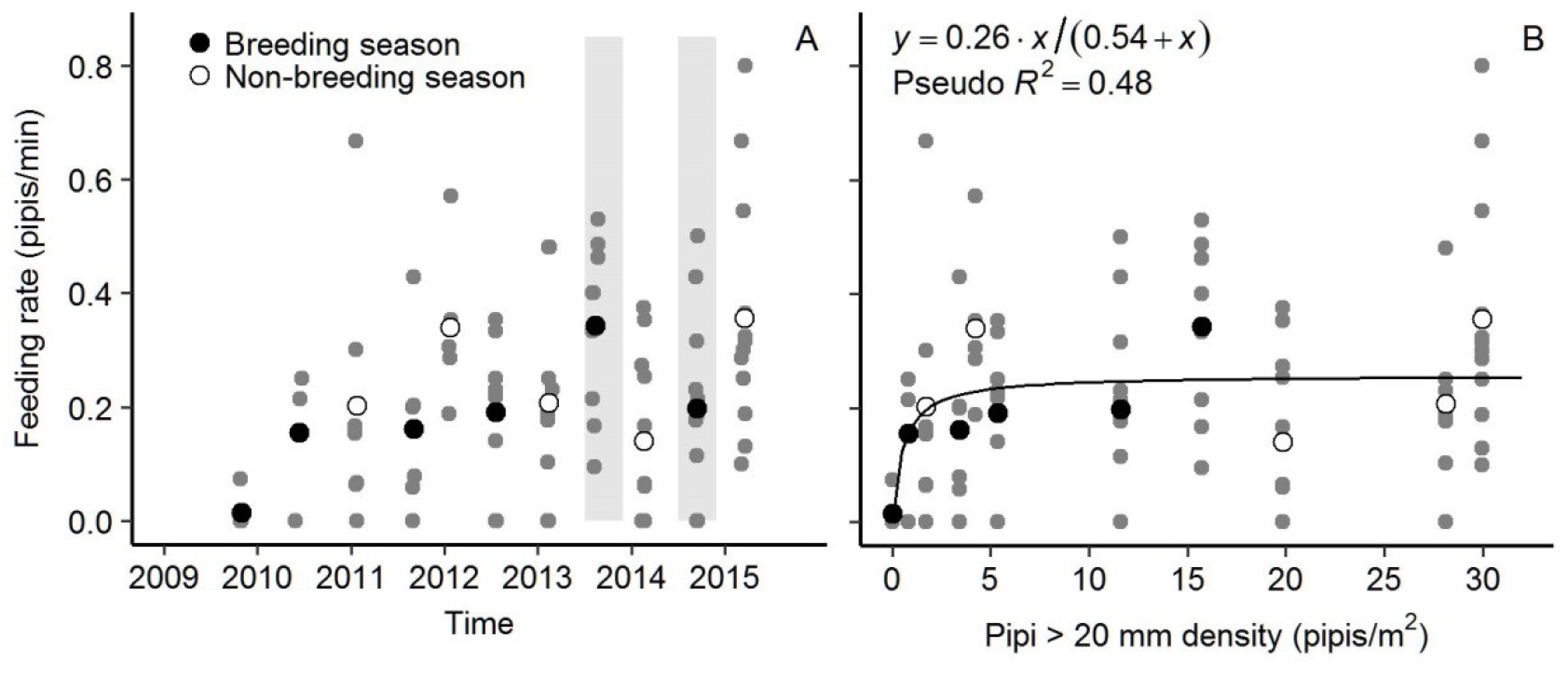
Oystercatcher pipi feeding rates at South Ballina Beach over time (A) and functional response to increasing pipi density (B). Black-filled circles are breeding season means and white-filled circles are non-breeding season means. The time scale (A) is continuous, with tick marks at 1 Jan. Light-grey vertical bands (A) indicate commercial shellfishing seasons. The asymptotic hyperbolic function (B) was fitted by non-linear least-squares regression. Pseudo-*R*^2^ is simply the squared Pearson’s product-moment linear correlation between observed and fitted values.

## Discussion

Predicted responses of Australian pied oystercatchers to increasing pipi abundance at South Ballina Beach were confirmed with the numerical response mainly driven by non-territorial oystercatchers, and feeding rates increasing to an asymptote. The link between numerical and functional responses for shorebirds is well known (Colwell 2010) and these findings from one site should be relevant for other sandy ocean beaches in NSW with pipis and oystercatchers.

### Diet and prey size selection

Length-frequency distributions for live pipis > 20 mm and shells left by feeding oystercatchers agreed closely over six sampling occasions. For comparison, Owner (1997) compared live pipi and shell samples from one location on South Ballina Beach where a flock of non-territorial oystercatchers had been feeding and those two length-frequency distributions agreed very closely for live pipi > 20 mm (*n* = 78 pipis, *m* = 200 shells, *χ*^2^ = 2.6, *df* = 9, *P* = 0.98; computed from histograms in Fig. 4.2 of Owner 1997). Taylor and Taylor (2005) studied oystercatchers taking the clam *Mactra rufescens* on a beach in Tasmania and again reported no difference in mean length between live clams and shells left by feeding oystercatchers. Taylor and Taylor’s (2005) proposed explanation was that profitability (biomass ÷ handling time) did not increase for clams from 40–62 mm although that result did not include smaller clams that were taken. The full range of clams taken must be covered in such studies because biomass increases exponentially with shell length (*e.g.* McLachlan & Hanekom 1979; Murray-Jones 1999 for *Donax* species). Taylor and Taylor’s (2005) alternative and more general explanation for no size selection among beach clams taken by oystercatchers was that oystercatchers were unable to distinguish between sizes of buried clams or that they had little time for careful size assessments when foraging in the swash zone. Pipis feed by extending their siphons to the sand-water interface and oystercatchers can locate pipis by visual searching for pipi siphons. It is not known if siphons give a reliable indication of the size of the animal and, anyhow, oystercatchers have little time to act before the next wave arrives.

The ‘tried and tested’ method for measuring prey size (and thereby intake rates) in Eurasian oystercatcher feeding studies involves visual estimation of prey size in comparison to bill length or other known-size marks (Goss-Custard *et al.* 1987). Goss-Custard *et al.* (2002) cautioned that shell collections may be biased towards larger shells, which are more likely to be seen and collected than are smaller shells. Another concern for sandy ocean beaches is size-related transport and deposition of shells by the waves. Any differences found between length-frequency distributions in shell collections compared to live prey samples are ambiguous: they could represent either prey size selection or method bias. Conversely, the absence of substantial differences observed in this study and in Owner (1997) suggests both small bias and a lack of pipi size selection.

Observational studies of shorebird diets are always incomplete because small prey cannot be identified. Small prey were 41% of prey eaten in this study and polychaete worms were 2%. Harrison (2009) sampled benthic macroinvertebrates on six beaches (*n* = 11 ‘sites’ × 3 transects × 3 quadrats, including five sites on South Ballina Beach) using a one mm sieve mesh. Pipis averaged 36% of total numerical density, tiny crustaceans (amphipods, isopods) < 6 mm averaged 38% and polychaete worms < 32 mm averaged 19% (although larger worms are present, are used by beach fishers for bait and occasionally taken by oystercatchers, pers. obs). Agreement between the 38–57% small prey available in Harrison (2009) and 41% small prey taken in this study suggests no size selection among different prey types for oystercatchers on sandy ocean beaches. Small prey < 6 mm must have small biomass relative to bivalves > 20 mm and can justifiably be ignored in oystercatcher feeding studies where large bivalves are frequently taken. Two prey categories are, therefore, proposed for oystercatchers on sandy ocean beaches: 1) small prey with high finding cost, low handling cost and low value, and 2) large prey (generally bivalves) with high finding cost, high handling cost and high value. The finding cost (walking, running, searching, sewing, probing) is high for both prey types because a large part of swash zone foraging time is spent avoiding waves. Once found, small prey are taken because the additional cost (*e.g.* peck and swallow) is small.

### Functional response

Oystercatcher feeding rates increased rapidly to an asymptote of c. 0.26 pipis per min. of active foraging time. Like many other shorebird functional responses reviewed in Goss-Custard *et al.* (2006), this asymptote was reached at low prey densities and feeding rates were noisy. Larger sample sizes and higher precision could be achieved by measuring pipi density in the swash zone together with each feeding observation and then averaging measurements within narrow density ranges. A related concern is whether beach scale sampling is valid. Functional response results in the literature are often extrapolated from the measurement scale (*e.g.* study plots) to the site scale (*e.g.* the estuary) when making inferences about the local oystercatcher population (or even the population; Goss-Custard 1996). This assumes that the functional response is scale invariant. This study directly measured the recovery in pipis and oystercatchers at the beach scale. Moreover, measurements of prey abundance are irrelevant at the functional response asymptote (Goss-Custard *et al.* 2006).

For comparison with results from this study, Owner (1997) previously measured feeding rates for one oystercatcher pair on South Ballina Beach in 1997 at 0.5–0.8 pipis/min of visual plus tactile searching time. Recalculating feeding rates from this study for the same time-base (searching plus sewing time) and refitting the asymptotic hyperbolic function gives an asymptote 0.9 pipis/min (95% CI 0.4–1.5). Thus, asymptotic feeding rates for pipi > 20 mm mean density up to 30 pipis/m^2^ in this study were similar to those at 107/m^2^ in Owner (1997; *n* = 5 transects). This 107/m^2^ density is adjusted for 18% pipis ≤ 20 mm (using length-frequency data in Fig 4.2 of Owner 1997) but an average of one pipi every 10 cm still seems unusually high.

Feeding observation sample sizes in this study were small, intake-rates were not measured and these results provide only a first approximation of the functional response for oystercatchers and pipis. Despite these limitations, mean feeding rates supported the asymptotic functional response model and clearly not the ‘more is better’ linear functional response model. More detailed studies of oystercatcher feeding are welcome but the evidence so far indicates no prey size selection among pipis > 20 mm and a functional response with a low half-asymptote density.

### Numerical response

Non-territorial oystercatchers responded more quickly to increasing pipi abundance than did territorial oystercatchers. Non-territorial oystercatchers are free to move along the coastline and, following Owner and Rohweder (2003), it is proposed that South Ballina Beach attracted more oystercatchers as pipi abundance increased. Non-territorial oystercatcher counts increased strongly from Jan 2013 onwards with corresponding pipi > 20 mm mean densities ≥ 11/m^2^. For comparison, Harrison (2009) reported a mean pipi density of 6/m^2^ for six beaches without oystercatchers and 26/m^2^ for eight beaches with oystercatchers. Seasonal variation in the final three years of this study further shows the fast response of non-territorial oystercatchers, however interpretation of this result is ambiguous. One explanation is that higher occupancy of territories in the breeding season reduced habitat available to non-territorial birds. Another is that non-territorial oystercatcher decreased with lower pipi abundance during the commercial pipi fishing seasons, however pipi abundance on South Ballina Beach was still greater than other beaches in northern NSW (unpub. data; Owner and Rohweder 2003).

The breeding season mean count of 43 adult-plumage oystercatchers in Sep 2014, at the end of this study, is similar to 38 total oystercatchers in 1997 (Owner & Rohweder 2003) and 38−42 adults in 2003–2005 (Harrison 2009) except that the population structures were different. In this study, the maximum number of territories was 11 (22 birds) with pipi > 20 mm mean density 12/m^2^ (increasing to 30/m^2^ on the final sampling occasion). Wellman (*et al.* 2000) reported 15 breeding pairs on South Ballina Beach (30 birds) in 1997 and Owner (1997) reported a mean pipi > 20 mm mean density of 107/m^2^. Harrison (2009) counted 18 territories (36 birds), 15 breeding pairs and mean pipi density 20/m^2^ (*n* = 5 sites × 3 transects × 3 quadrats) in 2003.

Eight oystercatcher territories were identified in the 2009 and early 2010 breeding seasons, even though pipi abundance was c. zero. Breeding season oystercatcher territories continued to increase slowly despite lower pipi abundance during the 2013 and 2014 commercial pipi fishing seasons. These results indicate that oystercatcher territories are not particularly sensitive to pipi abundance.

Like other oystercatcher species, the density of Australian pied oystercatcher breeding pairs is usually limited by territorial behaviour (Taylor *et al.* 2014). Competition for territories is strong and vacancies are rapidly filled by birds from the non-breeding surplus (*e.g.*Harrison 2009). One proposed explanation for the slow response in territorial counts in this study is that movements of oystercatchers out of South Ballina Beach during the 2003–2009 pipi crash disrupted the local social order. When pipi abundance was declining, there was an exodus of non-territorial birds first, followed by a decline in breeding pairs (Harrison 2009). The asymptotic functional response predicts low feeding rates at pipi densities near zero and the proposed explanation for those exoduses is food shortages (Harrison 2009). When pipi abundance increased, non-territorial birds moved in but did not immediately take up territories. These newcomers are assumed to start near the bottom of the local social order and are not expected to occupy and defend vacant breeding space (Ens *et al.* 1996c). In the longer term, there are other processes that could be important, in particular breeding habitat degradation and loss to human recreation and development (see Taylor *et al.* 2014 for a review). Numerical response models from this study do show an association between oystercatcher counts and pipi abundance but they do not predict population changes into the future.

### Pipi harvesting

It is not known what drives ‘pipi crashes’. The overfishing hypothesis (*e.g.*Owner and Rohweder 2003; Harrison 2009) proposes that fishing can drive stocks to near zero. This can occur because of the phenomenon of ‘hyperstability’ for species such as pipis that form dense and easily harvested aggregations, where catch rates can remain high despite large decreases in abundance (Harley *et al.* 2001). The recruitment variability hypothesis proposes that weak or zero recruitment can drive declines. Murray-Jones (1999) suggested that weak or zero pipi recruitment can result from post-settlement mortality. For *Katelysia scalarina* clams in Tasmanian estuaries, Taylor (2004) suggested that the clam harvesting method, which disturbed the sediment, resulted in recruitment failure.

Fishery catch data are considered unreliable for measuring pipi population dynamics (Ferguson *et al.* 2015), not publicly available, and there are no continuous fishery-independent data before 2009 to evaluate the pipi overharvesting hypothesis. There is some anecdotal evidence for the recruitment variability hypothesis. Pipi harvesting on South Ballina Beach stopped in 2004 (Harrison 2009) and yet pipi abundance continued to decline. In 2003, Harrison’s (2009) mean pipi density was 21/m^2^ for four beaches in northern NSW with breeding oystercatchers, including 20/m^2^ at South Ballina. In 2005, mean pipi density was 6/m^2^ (2005 results for individual beaches were not reported). In 2009, pipi abundance on South Ballina Beach was c. zero (this study). Mass mortality of pipi recruits (*i.e.* recruitment failure) was incidentally observed on South Ballina Beach in Nov 2007, during the 2003–2009 pipi crash (B. Moffatt, pers. comm. 2015). During the recent pipi crash and throughout this study, there are no known reports of mass mortality events involving adult pipis. Strong recruitment events in Jul 2012 and Sep 2014 supported commercial fishing and large increases in non-territorial oystercatcher counts on South Ballina Beach. Unlike Taylor (2004), simultaneous commercial shellfishing did not prevent pipi recruitment. Pipis are fast-growing and short-lived and it is proposed that the sustainability of pipi stocks requires frequent strong recruitment events.

Like Gray (2016a,2016b) BACI analyses failed to detect fishing impacts. More precise measurements require larger sample sizes (*i.e.* numbers of transects) and, to minimise the confounding effects of rapid growth for small pipis, sampling should be performed just before the start of the fishing season and soon after the peak in fishing activity. Even then, the BACI design is difficult to apply. Observed fishing-associated declines were temporary. Fishery management zones and spatial closures are not randomly designed and pipi populations can vary among and within beaches for factors other than fishing including beach morphodynamics (Short 2007) and settlement of recruits (Murray-Jones & Ayr 1997). Observations from the 2003–2009 pipi crash and subsequent recovery suggest that a longer-term study of fishing impacts is required.

## Acknowledgements

Eight reviewers have given feedback on successive versions of this paper. Reasons for this paper being unpublished are (in no particular order): 1) it concerns only one study site, 2) few researchers have any experience of oystercatchers on sandy ocean beaches, 3) field methods were different to or less detailed than established, European methods, and 4) many oystercatcher conservationists are against shellfishing whereas this paper is not.

